# Role of ribosomal protein bS1 in orthogonal mRNA start codon selection

**DOI:** 10.1101/2024.10.14.618353

**Authors:** Kristina V. Boyko, Rebecca A. Bernstein, Minji Kim, Jamie H. D. Cate

## Abstract

In many bacteria, the location of the mRNA start codon is determined by a short ribosome binding site sequence that base pairs with the 3′-end of 16S ribosomal RNA (rRNA) in the 30S subunit. Many groups have changed these short sequences, termed the Shine-Dalgarno (SD) sequence in the mRNA and the anti-Shine-Dalgarno (ASD) sequence in 16S rRNA, to create “orthogonal” ribosomes to enable the synthesis of orthogonal polymers in the presence of the endogenous translation machinery. However, orthogonal ribosomes are prone to SD-independent translation. Ribosomal protein bS1, which binds to the 30S ribosomal subunit, is thought to promote translation initiation by shuttling mRNA to the ribosome. Thus, a better understanding of how the SD and bS1 contribute to start codon selection could help efforts to improve the orthogonality of ribosomes. Here we engineered the *Escherichia coli* ribosome to prevent binding of bS1 to the 30S subunit, to separate the activity of bS1 binding to the ribosome from the role of the mRNA SD sequence in start codon selection. We find that ribosomes lacking bS1 are slightly less active than wild-type ribosomes *in vitro*. Furthermore, orthogonal 30S subunits lacking bS1 do not have improved orthogonality. Our findings suggest that mRNA features outside the SD sequence and independent of bS1 binding to the ribosome likely contribute to start codon selection and the lack of orthogonality of present orthogonal ribosomes.

## INTRODUCTION

Translation initiation in bacteria involves the formation of a 30S initiation complex, in which the start codon of the mRNA pairs with the initiator fMet-tRNA anticodon in the P site of the small (30S) ribosomal subunit. The positioning of the start codon in the 30S initiation complex is aided by the SD sequence 5′ of the start codon, which base pairs to the complementary ASD sequence in 16S rRNA. Characteristics of the SD sequence, including its presence or absence, spacer length from the SD sequence to the start codon, or SD sequence composition, can determine translation efficiency.^1^ For example, many bacteria encode leaderless mRNAs, or mRNAs that do not contain an identifiable SD sequence, and only 54% of *E. coli* genes have an SD sequence.^2^ RNA-seq analysis has also revealed that leaderless mRNAs can comprise up to 70% of transcripts in other bacteria as well as archaea.^3^ In mRNAs possessing an SD sequence, the number of nucleotides between the start codon and the SD sequence affects the levels of protein expression.^4^ Finally, nucleotides which compose the SD sequence can also affect translation. For example, in *E. coli* the canonical SD sequence (5′-UAAGGAGG-3′) yields better translation efficiency than a 5′-AAGGA-3′ sequence.^5^ Taken together, these findings suggest that the role of the SD sequence in the mechanism of translation initiation does not follow a simple set of rules, and can vary widely across bacteria.

Mechanisms of gene expression in *E. coli* have been the most widely studied in bacteria, making this model organism the most popular host for synthetic biology applications. These include efforts to expand the genetic code to enable noncanonical amino acid incorporation into proteins for the synthesis of non-proteinogenic polymers by the ribosome.^6^ To enable genetic code expansion, many groups have engineered cells with an orthogonal ribosome, a secondary ribosome to translate orthogonal mRNAs independent of the endogenous translation machinery, along with orthogonal mRNAs (omRNAs).^7^ The orthogonal ribosome can be engineered to incorporate non-canonical amino acids while the endogenous ribosome can continue translating cellular mRNAs, preventing cellular growth defects.^8,9^ A central aspect of engineering mRNAs encoding non-canonical proteins to be orthogonal to the endogenous translation machinery is to include an orthogonal ribosome binding site (RBS) on the 16S rRNA and an orthogonal SD (oSD) sequence on the mRNA, for start codon selection.^10,11,7^ To ensure that the orthogonal ribosome only translates these omRNAs, 16S rRNA in the orthogonal 30S subunit (o30S) harbors an orthogonal ASD (oASD) sequence that base pairs with the omRNA RBS. This combination of oSD in the omRNA and oASD in the o30S subunit, in theory, should ensure that the target omRNA is only translated by the orthogonal ribosome, while the the orthogonal ribosome does not translate endogenous mRNAs.

Multiple examples of orthogonal RBS have been reported, in which both the 16S rRNA and mRNA RBS sequences have been engineered. Approaches to identify these sequences involved positive and negative selections as well as computational methods.^7,12^ However, recent studies in *E. coli* have shown that the SD sequence does not fully dictate whether or not an mRNA transcript is translated. Single-molecule tracking techniques in live cells revealed that translating ribosomes with oASD sequences in their 16S rRNA bind to endogenous mRNAs.^13^ This finding is further supported by ribosome profiling experiments that reveal that different ASD sequences in 16S rRNA do not prevent orthogonal ribosomes from translating cellular mRNAs.^14^ Taken together, this cross-reactivity of orthogonal ribosomes with endogenous mRNAs limits the current utility of these orthogonal systems (Figure 1A).

**Figure 1.**
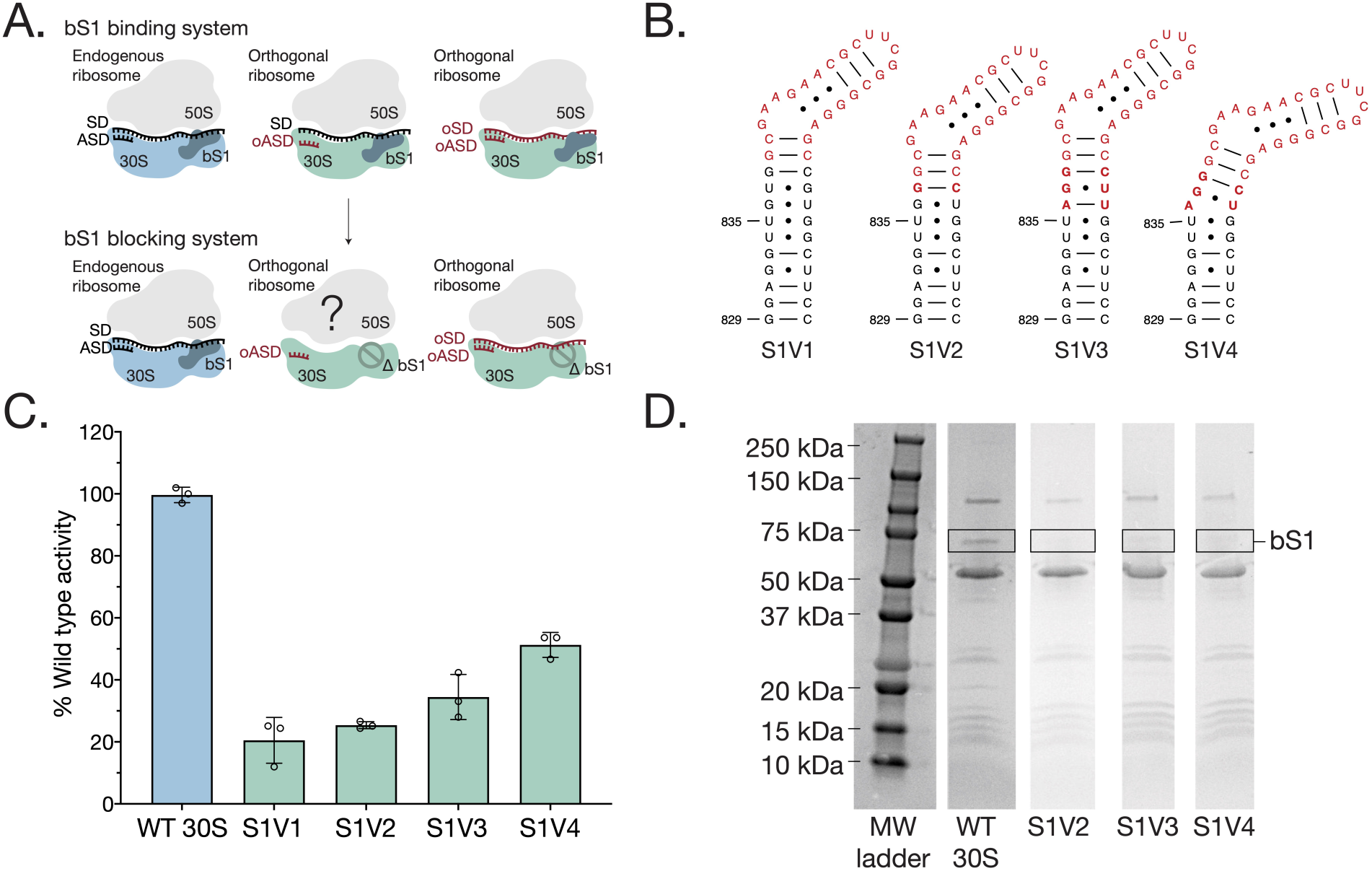
Model and characterization of bS1 blocking ribosomes. **A.** Proposed model of orthogonal ribosomes in a bS1 binding system and a bS1 blocking system. Current orthogonal ribosomes that undergo SD independent translation translate orthogonal mRNA (right) as well as endogenous mRNA (middle), possibly due to bS1. A bS1 blocking system would prevent binding of bS1 to orthogonal ribosomes, which could prevent translation of endogenous mRNA (middle) while encouraging translation of orthogonal mRNA (right). Endogenous mRNA is shown in black, and orthogonal mRNA in red. **B.** Secondary structure of all versions of the bS1 blocking mutants inserted into helix h26 of *E. coli* 16S rRNA. Nucleotides colored red are added bases. Nucleotides in bold highlight differences between versions S1V1-S1V4. **C.** *In vitro* activity assay of wild-type 30S and bS1 blocking mutants with wild-type 50S subunits. Activity corresponds to luminescence of synthesized nanoluciferase, normalized to wild-type ribosomes. Error bars indicate standard deviation for n=3 replicates. **D.** 10% SDS PAGE gel of wild-type 30S subunits and versions S1V2-S1V4 of the bS1 blocking mutants. The black box highlights bS1. The relative intensity of the maltose and MS2 binding fusion protein used for purification, as well as the other ribosomal proteins, is consistent across all lanes, indicating that the same amount of protein was loaded for each sample.

In *E. coli*, ribosomal protein bS1 is thought to contribute to translation initiation using mechanisms independent of the SD sequence.^15^ Ribosomal protein bS1 is composed of multiple RNA-binding domains and is, in some cases, thought of as an additional translation factor due to its reversible binding to the ribosome.^15^ Protein bS1 also has high affinity for mRNA and is thought to facilitate translation for highly structured mRNAs.^16^ However, the translation of leaderless mRNAs, as well as of mRNAs that have an RBS close to the 5′ end, is independent of bS1.^16^ Recent cryo-EM structures of *E. coli* ribosomes with truncated ribosomal protein bS1 reveal that bS1 binds to mRNA, just upstream of the SD sequence.^17^ In addition to binding to the mRNA, bS1 simultaneously binds to ribosomal protein uS2 adjacent to the SD-ASD interaction site, as well as to 16S in this region.^18^ Taken together, these mechanistic and structural findings have led to the model that bS1 acts to recruit mRNA to the ribosome in an SD independent manner.

We reasoned that the ability of ribosomal protein bS1 to recruit mRNAs to the ribosome in an SD-independent manner might be responsible for the lack of orthogonality seen with present orthogonal ribosomes and omRNAs. We therefore tested whether orthogonal ribosomes could be engineered to rely more on SD and ASD binding, by engineering an orthogonal 16S rRNA that sterically prevents bS1 binding to the 30S ribosomal subunit. In this case, ribosomal protein bS1, an essential protein in *E. coli*, would continue to bind to the endogenous ribosome and translate endogenous mRNAs.^15^ However, the orthogonal ribosomes that have an orthogonal ASD and translate omRNAs, would not bind bS1 and eliminate this aspect of bS1’s contribution to mRNA recruitment to the ribosome (Figure 1A). Using cryo-EM and *in vitro* and cell-based translation assays, we found that preventing bS1 binding to the 30S subunit does not affect ribosome orthogonality *in vitro* or *in vivo*, indicating that SD-independent mRNA translation is not solely dependent on bS1 binding to the ribosome.

## MATERIALS AND METHODS

### Plasmid preparation

A pLK35 plasmid with an MS2 tagged located in 16S rRNA helix h6 was a gift from the Schepartz lab at UC Berkeley.^19^ The helix h26 extensions were introduced using the In-Fusion Cloning Kit (Takara Bio) with the primers listed below. The oASD was also cloned into the wild type plasmid as well as S1V4 using the same method and the following primers:

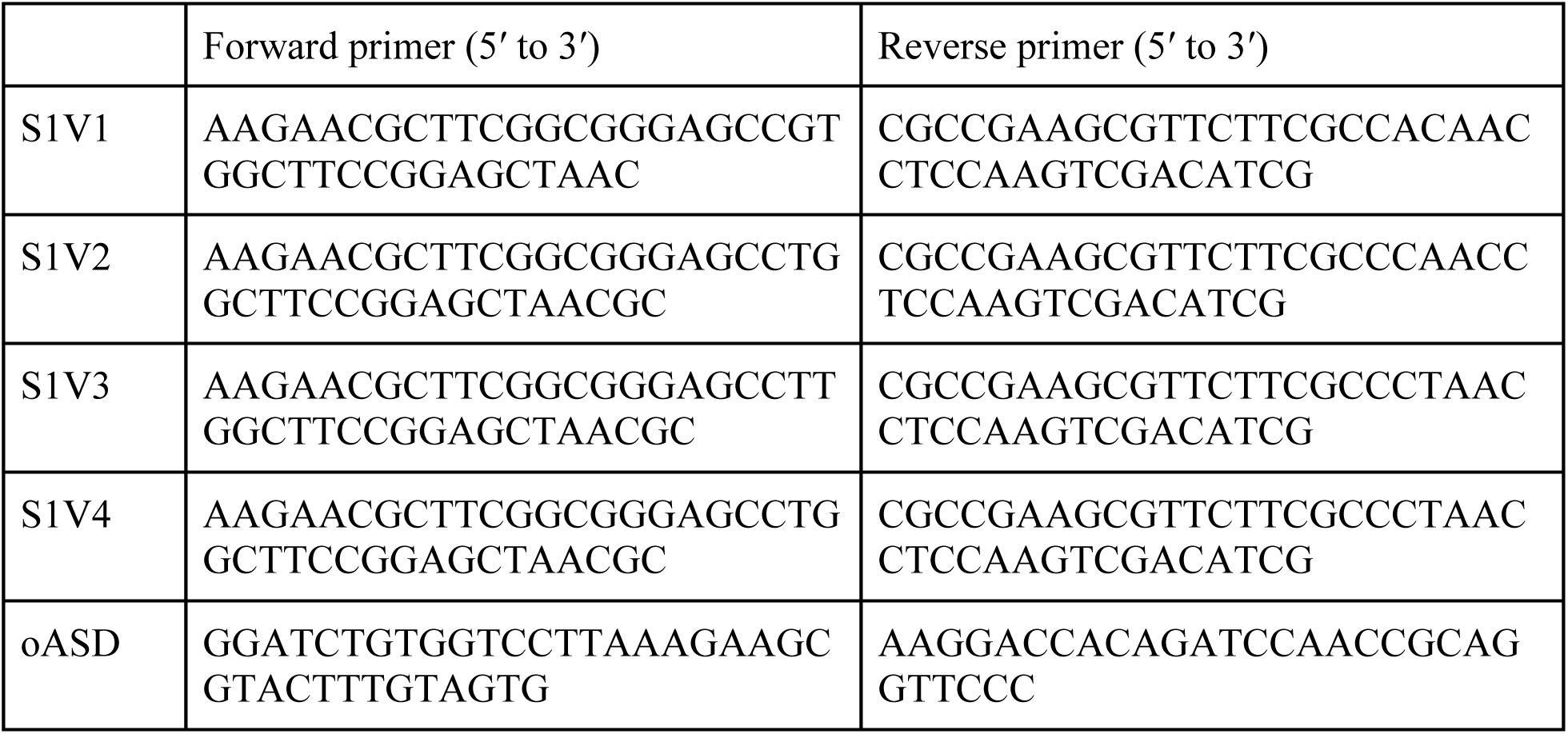

### Reporter constructs

An oSD was cloned into a nanoluciferase reporter using the In-Fusion Cloning Kit (Takara Bio) and the primers listed below. A GFP gene block with overhangs (Twist) was cloned into plasmid pJH474_pTech_Para-supP (Addgene) using the In-Fusion Cloning Kit.^20^ An oSD was cloned into the GFP reporter using the primers listed below.

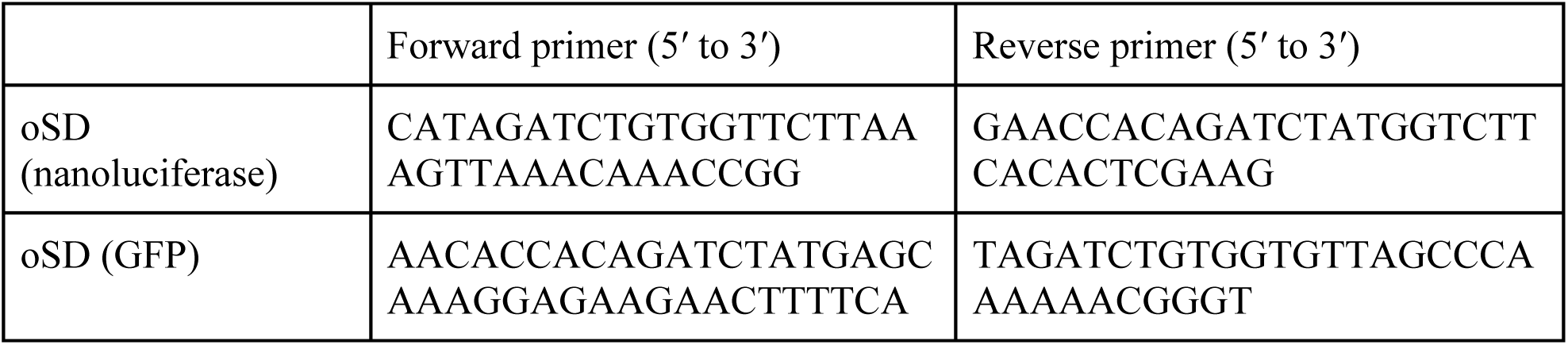

### MBP-MS2 protein expression and purification

A pMAL-c2 plasmid (NEB) encoding an N-terminally 6xHis tagged MBP-MS2 fusion protein was transformed into *E. coli* BL21 cells.^21^ A colony from the transformation was grown overnight in 10 mL of LB broth containing 100 μg/mL ampicillin. The culture was transferred to 1 L of ZYM-5052 autoinduction media^22^ with 100 μg/mL ampicillin and grown overnight at 37 ℃ with shaking at 175 RPM. Cells were pelleted by centrifugation (6000 xg, 4 ℃, 20 minutes), resuspended in lysis buffer (20 mM HEPES pH 7.5, 250 mM KCl, 10 mM imidazole), and then lysed using sonication. The resulting lysate was clarified by centrifugation (33746 xg, 4 ℃, 45 minutes) and subsequently filtered through a 0.2 μm filter. The lysate was loaded onto a gravity column with 3 mL of Cobalt Resin (Thermo Scientific). The column was washed four times with three column volumes of lysis buffer. The protein was eluted into 1 mL fractions using an elution buffer (20 mM HEPES pH 7.5, 250 mM KCl, 2 mM BME, 500 mM imidazole). The purified protein was buffer exchanged to MS2-150 Buffer (20 mM HEPES pH 7.5, 150 mM KCl, 1 mM EDTA) and concentrated to 10 mg/mL using 30 kDa spin filters.

### Mutant ribosome expression and purification

Plasmids containing the 16S rRNA variants designed to block bS1 binding were transformed into *E. coli* Mach1 competent cells (Thermo Fisher) and cultures were grown overnight in 10 mL of LB broth containing 100 μg/mL ampicillin. Cultures were transferred to 1 L LB containing 100 μg/mL ampicillin and incubated at 37 ℃ with shaking at 175 RPM. Cultures were induced with Isopropyl β-d-1-thiogalactopyranoside (IPTG) at OD_600_ = 0.4, with a final IPTG concentration of 0.5 mM, for 4 hours. Cells were pelleted, resuspended in ribosome buffer A-1 (20 mM Tris HCl pH 7.5, 100 mM NH_4_Cl, 1 mM MgCl_2_, 0.5 mM EDTA), and then lysed using sonication. The resulting lysate was clarified by centrifugation (33746 xg, 4 ℃, 45 minutes) and loaded into Ti-45 tubes. A sucrose cushion was prepared by adding 24 mL buffer B (20 mM Tris HCl pH 7.5, 0.5 mM EDTA, 100 mM NH_4_Cl, 10 mM MgCl_2_, 0.5 M sucrose) and 17 mL buffer C (20 mM Tris HCl pH 7.5, 0.5 mM EDTA, 60 mM NH_4_Cl, 6 mM MgCl_2_, 0.706 M sucrose) to the Ti-45 tubes and the ribosomes were pelleted by centrifugation (27000 RPM or 57,000 xg, 4 ℃, 15 hours). Crude ribosomes were resuspended in ribosome buffer A-1, clarified by centrifugation (21130 xg, 4 ℃, 25 minutes), and subsequently filtered using a 0.2 μm filter.

Crude ribosomes (>60 mg) were diluted to 15 mg/mL in ribosome buffer A-1, and purified at 4 ℃. A 5 mL MBP-trap column (Cytiva) was buffer exchanged to MS2-150 buffer and 10 mg of MBP-MS2 was diluted to 3 mL in MS2-150 buffer before being loaded onto the column. The column was washed with 5 column volumes of ribosome buffer A-1 and crude ribosomes were then slowly loaded onto the column. The column was washed with another 5 column volumes of ribosome buffer A-1 and then 10 column volumes of ribosome buffer A-250 (20 mM Tris HCl pH 7.5, 250 mM NH_4_Cl, 1 mM MgCl_2_, 0.5 mM EDTA). Ribosomes were eluted into 1 mL fractions using an elution buffer containing 10 mM maltose (20 mM Tris HCl pH 7.5, 100 mM NH_4_Cl, 1 mM MgCl_2_, 0.5 mM EDTA, 10 mM maltose). The purified ribosomes were buffer exchanged to ribosome buffer A-1, concentrated to 3000 nM using 100 kDa spin filters (using the approximation of 1 A_260_ = 72 nM), and flash frozen in liquid nitrogen.

### Wild-type ribosome expression and purification

Wild-type 30S and 50S subunits were expressed and purified as previously described.^23^ Briefly, *E. coli* MRE600 cells were grown for four hours before being pelleted and resuspended in ribosome buffer A-1 and subsequently sonicated. The lysate was clarified by centrifugation and crude ribosomes were isolated using a sucrose cushion at 4 ℃. The crude ribosomes were then separated on a 20-40% sucrose gradient using a dissociation buffer (20 mM Tris-HCl pH 7.5, 60 mM NH4Cl, 1 mM MgCl_2_, 5 mM EDTA) and individual subunits were isolated via fractionation. The subunits were buffer exchanged to ribosome buffer A-1 for storage.

### Ribosome purity assay

60 μg of purified mutant ribosomes were heated at 95 ℃ for 5 minutes. Lithium chloride precipitation solution (Invitrogen) was then added at 2X the volume of ribosomes on ice and the samples were incubated at -20 ℃ for one hour to precipitate the rRNA. The rRNA was pelleted by centrifugation (21130 xg, 4 ℃, 25 minutes) and the supernatant was vacuum aspirated. The rRNA was washed with 40 μL 70% ethanol, pelleted by centrifugation (21130 xg, 4 ℃, 5 minutes), and the supernatant was vacuum aspirated. The resulting rRNA was resuspended in RNase-free water. Primers flanking the MS2 tag in the 16S rRNA, shown below, were used in a reverse transcription reaction using the OneStep RT-PCR kit (Qiagen). The resulting cDNA was resolved on a 10% TBE-gel and visualized with SYBR Safe DNA Gel Stain (Supplementary Figure 1). For RT-PCR, 1 mL of cells were spun down after four hours of induction and lysed via a 5 minute heat shock at 98 ℃. The debris was pelleted and the RT-PCR was performed as described above.

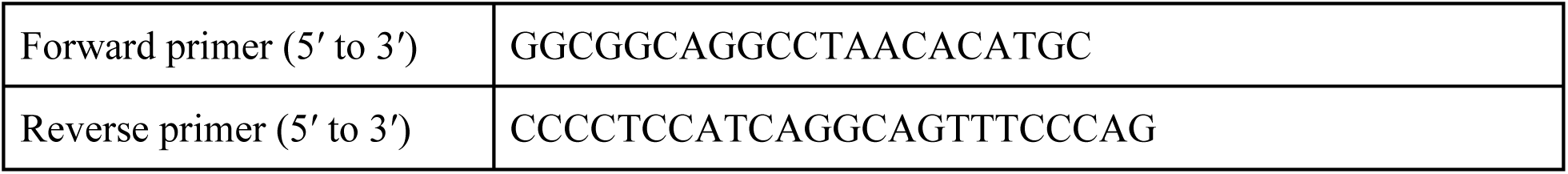

### Reassociation sucrose gradients

20-40% sucrose gradients (20 mM Tris-HCl pH 7.5, 60 mM NH_4_Cl, 10 mM MgCl_2_, 0.5 mM EDTA) were prepared in SW 41 Ti tubes. 500 nM 50S and 1000 nM 30S were incubated at 37 ℃ for 45 minutes before being added to the gradients (20 mM Tris-HCl pH 7.5, 60 mM NH_4_Cl, 10 mM MgCl_2_, 0.5 mM EDTA). The gradients were run overnight at 4 ℃ in an SW 41 Ti swinging bucket rotor (Beckman Coulter) and subject to 5.45×10^11^ ω^2^t total centrifugal force (27,000 rpm, 16 hours). The gradients were analyzed on a Biocomp fractionator.

### Cryo-EM sample preparation

70S ribosome fractions were collected from a 20-40% sucrose gradient using a Biocomp fractionator and buffer exchanged to cryo-EM buffer (20 mM HEPES pH 7.5, 10 mM Mg(OAc)_2_, 2 mM DTT, 100 mM KCl). The sample was incubated at 37 ℃ for 45 minutes before use. 300 mesh 100 R 1.2/1.3 UltraAuFoil grids (Quantifoil) with 2 nm carbon were glow discharged in a PELCO easiGlow for 12 seconds at 25 mAmp and 0.37 mBar before 4 μL of sample were added to each grid. The samples were incubated on the grid for one minute before being washed three times in 100 μL of cryo-EM buffer. The grids were blotted on a FEI Mark IV Vitrobot using a blot force of 6 and a blot time of 3 s at 4 ℃ in 100% humidity. The grids were then plunged in liquid ethane.

### Cryo-EM data acquisition and processing workflow

Movies were collected on a 300 kV Titan Krios microscope with a BIO-energy filter and Gatan K3 camera as previously described,^24^ using a super-resolution pixel size of 0.41 Å and a physical pixel size of 0.82 Å. SerialEM^25^ was used for automated data collection over a defocus range of -0.5 to -1.5 µm with an electron dose of 40 e^-^/Å^2^ over 40 frames.

Raw movies were imported into CryoSPARC 4.^30^ The movies were motion corrected using Patch Motion Correction^26^ before CTFFind4^27^ was used to estimate the CTFs of micrographs. Particles were picked using Template Picker with a 70S ribosome 2D template. Particles were extracted and Fourier-cropped to ⅛ of the box size. 2D classification was run with 100 classes and 32 classes that were consistent with 70S ribosomes were selected for a second round of 2D classification. The particles were re-extracted and Fourier-cropped to ¼ of the box size and particles were subjected to heterogeneous refinement using a 70S ribosome map from Watson et al.^24^ Classes consistent with 70S ribosomes were re-extracted at the full box size and then subjected to homogeneous refinement (Supplementary Figure 2). For the S1V3 and S1V4 maps, local refinement was done with a mask over the 30S subunit. The pixel sizes for the maps were calibrated using a simulated map from PDB 7K00. For modeling, the coordinates from the 30S subunit in 7K00 were used as an initial model. Real-space refinement in PHENIX was used to refine the model.^28^ Local resolution was determined using Local Resolution Estimation in Relion.^29^ Figures were prepared in ChimeraX.^30^

### *In vitro* translation reactions

An *in vitro* translation reaction using a ΔRibosome PURExpress kit (NEB) was prepared in a total volume of 10 μL containing the following components: 2 µL solution A, 0.63 µL factor mix, 0.13 µL RNase inhibitor, and 25 ng/µL nanoluciferase DNA template. The reaction also included 250 nM 50S ribosomal subunits and 500 nM 30S ribosomal subunits. The remaining volume consisted of Milli-Q water. The reaction was incubated at 37 ℃ for 45 minutes. 2 μL of the reaction product was combined with 30 μL of a 1:50 dilution of Nano-Glo substrate (Promega) into a 364-well microplate, and luminescence was detected with the Tecan Spark Microplate Reader.

### *In vivo* translation reactions

*E. coli* NEBExpress cells were co-transformed with two plasmids using heat shock. One plasmid contained an aminoglycoside phosphotransferase gene for kanamycin resistance and expressed superfolder GFP under control of an arabinose inducible promoter. The second plasmid harbored a beta-lactamase gene for ampicillin resistance and expressed an rrnB ribosomal RNA operon with MS2-tagged 16S rRNA. Cells were grown overnight in 1 mL of LB containing 100 μg/mL ampicillin and 100 μg/mL kanamycin. 100 μL of the overnight culture was transferred to 3 mL LB containing 100 μg/mL ampicillin and 100 μg/mL kanamycin and allowed to grow until they reached a cell density of OD_600_ = 0.2. Cultures were then induced with 0.5 mM IPTG and 0.1% arabinose final concentrations. Cultures were grown for 4 hours at 37 ℃ with shaking at 175 RPM. Cells were then pelleted and resuspended in 250 μL 1x PBS (pH 7.2), which was transferred to a clear 96-well microplate. GFP fluorescence and the OD_600_ were measured with the Tecan Spark Microplate Reader. GFP fluorescence was normalized to the OD_600_ optical density.

## RESULTS

### An RNA extension blocks bS1 from binding ribosomes

Previous structural studies have shown that bS1 binds to the 30S subunit near helix h26 in 16S rRNA.^17,18^ Given that helix h26 contains variable regions that are surface exposed and are not phylogenetically conserved,^31^ we introduced an rRNA extension to helix h26 between positions U835 and G851 to attempt to sterically block bS1 binding to the mutant ribosomes.^32^ However, we kept the conserved loop of the helix, which contains a stable UUCG tetraloop, constant (Figure 1B).^33^ We tested four different variable sequences that form kink-turn (k-turn) motifs to identify an RNA structural element that could sterically prevent bS1 from binding the 30S subunit (S1V1-S1V4).^34^

We first inserted a k-turn derived from an engineered RNA that introduced 14 additional bases into helix h26 in 16S rRNA (S1V1).^35^ However, the S1V1 30S subunit was only 20% active compared to tagged wild-type 30S subunits, as shown by *in vitro* translation of a nanoluciferase reporter (Figure 1C). We therefore tried introducing G-C base pairs at the base of the helix h26 extension to both stabilize and extend the helix, to create mutant S1V2. To test for bS1 depletion from the S1V2 30S subunit, we expressed and purified ribosomes via an MS2 hairpin on the 16S rRNA.^19^ After purification, we isolated and resolved the proteins from the samples on a 15% SDS-PAGE protein gel. As a control, we also tagged wild-type 16S rRNA and purified wild-type 30S subunits adjacent to the mutant ribosomes. The gel shows a band for bS1 in the wild-type 30S at the expected molecular weight of 61.2 kDa, consistent with previous studies.^36^ However this band is missing in the S1V2 mutant sample (Figure 1D). While S1V2 successfully prevented bS1 from binding the 30S subunit, this mutant was only 25% as active as wild type 30S subunits (Figure 1C). To search for more active 30S subunits, we hypothesized that we could further extend the helix to prevent bS1 binding and introduced an additional A-U pair at the base of the helix (S1V3, Figure 1B). The S1V3 mutation successfully disrupted bS1 binding to the 30S subunit (Figure 1D) and was about 35% as active as wild-type ribosomes (Figure 1C).

We determined a cryo-EM structure of a 70S ribosome with the S1V3 30S subunit and found that ribosomal protein uS2 was missing from the 30S subunit (Figure 2A, Supplementary Figure 3). This suggested that the A-U base pair extension may have extended and rotated the k-turn into a position that collides with uS2. To mitigate this potential collision, we removed the uridine from the A-U base pair in mutant S1V4, such that stacking of the A in the RNA helix would angle the helix away from ribosomal protein uS2. This final variant, S1V4, also successfully blocked bS1 from binding the 30S subunit (Figure 1D), and was about 50% as active as wild-type 30S subunits. S1V4 also readily formed 70S ribosomes with wild-type 50S subunits as shown by a reassociation sucrose gradient (Supplementary Figure 4). To ensure the S1V4 30S subunit maintained structural integrity, i.e. with respect to ribosomal protein uS2, we determined the cryo-EM structure of S1V4 to a global resolution of 1.8 Å and a local resolution of 4.4 Å around the h26 extension (Figure 2B, Supplementary Figure 5, Supplementary Figure 6). The structure reveals well resolved density for the base of helix h26. However, as the helix continues into the k-turn the density becomes weaker, indicating a flexible k-turn extension (Figure 2C). Cryo-EM density is lacking for ribosomal protein bS1 but is present for uS2 indicating that the extension successfully prevents bS1 binding while not disturbing binding of uS2 to the ribosome (Figure 2B).

**Figure 2.**
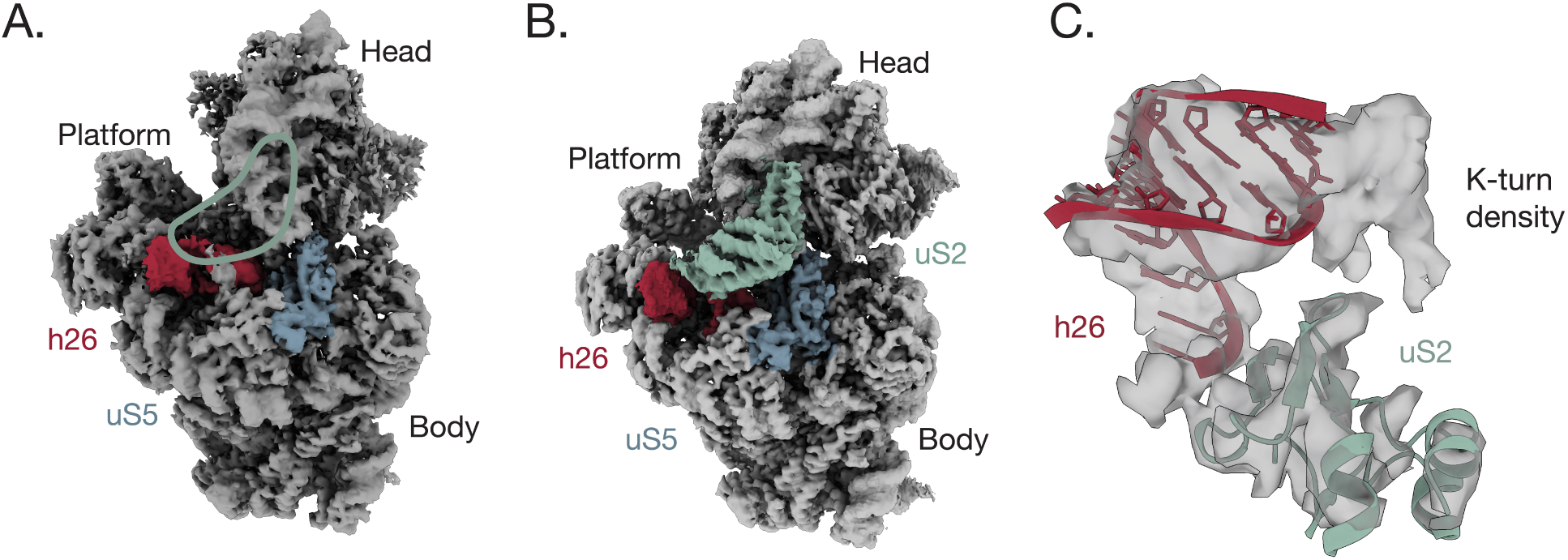
Structure of S1 blocking ribosomes. **A.** Cryo-EM density of 30S subunit S1V3, with density for ribosomal protein uS2 missing. Density is clearly resolved for protein uS5 and rRNA helix h26 (blue and red, EMD 47168). The green outline indicates where uS2 density would be. **B.** Cryo-EM density of 30S subunit S1V4. Density for uS2 is indicated in green and is well resolved. Density of uS5 is shown in blue and rRNA helix h26 is shown in red (EMD 47169). **C.** Cryo-EM density of the h26 k-turn extension (unmodeled) and uS2, in 30S subunit S1V4. The cryo-EM maps was blurred with a B-factor of 61 Å^2^.

### Ribosomal protein bS1 does not affect ribosome orthogonality

To test our hypothesis that orthogonal ribosomes can be made more dependent on the SD-ASD interaction by preventing bS1 binding to the 30S subunit, we used a previously-developed oASD sequence in both the wild-type 16S and S1V4 16S rRNA backgrounds.^12^ We also cloned the corresponding oSD sequence into a nanoluciferase reporter (Figure 3). Using a PURE *in vitro* transcription and translation (IVTT) system supplemented with purified 30S and 50S subunits, we measured nanoluciferase luminescence from IVTT reactions containing wild-type or S1V4 30S subunits, and mRNAs in all four possible combinations of wild-type SD and ASD, and oSD and oASD pairings. In the case of the wild-type 30S subunit, translation of the wild-type SD reporter is robust and the translation of the orthogonal SD-ASD pair is about 50% of wild-type levels (Figure 3). Consistent with published results, there is no detectable translation for non-cognate SD-ASD pairs when using wild-type 30S subunits capable of binding bS1.^37^ We observed similar results with S1V4, although there is slight cross-reactivity between the wild-type ASD and oSD. These results suggest that the SD-ASD interaction can be made highly orthogonal *in vitro*, and prevention of bS1 binding to the 30S subunit is inconsequential and possibly deleterious to orthogonality.

**Figure 3.**
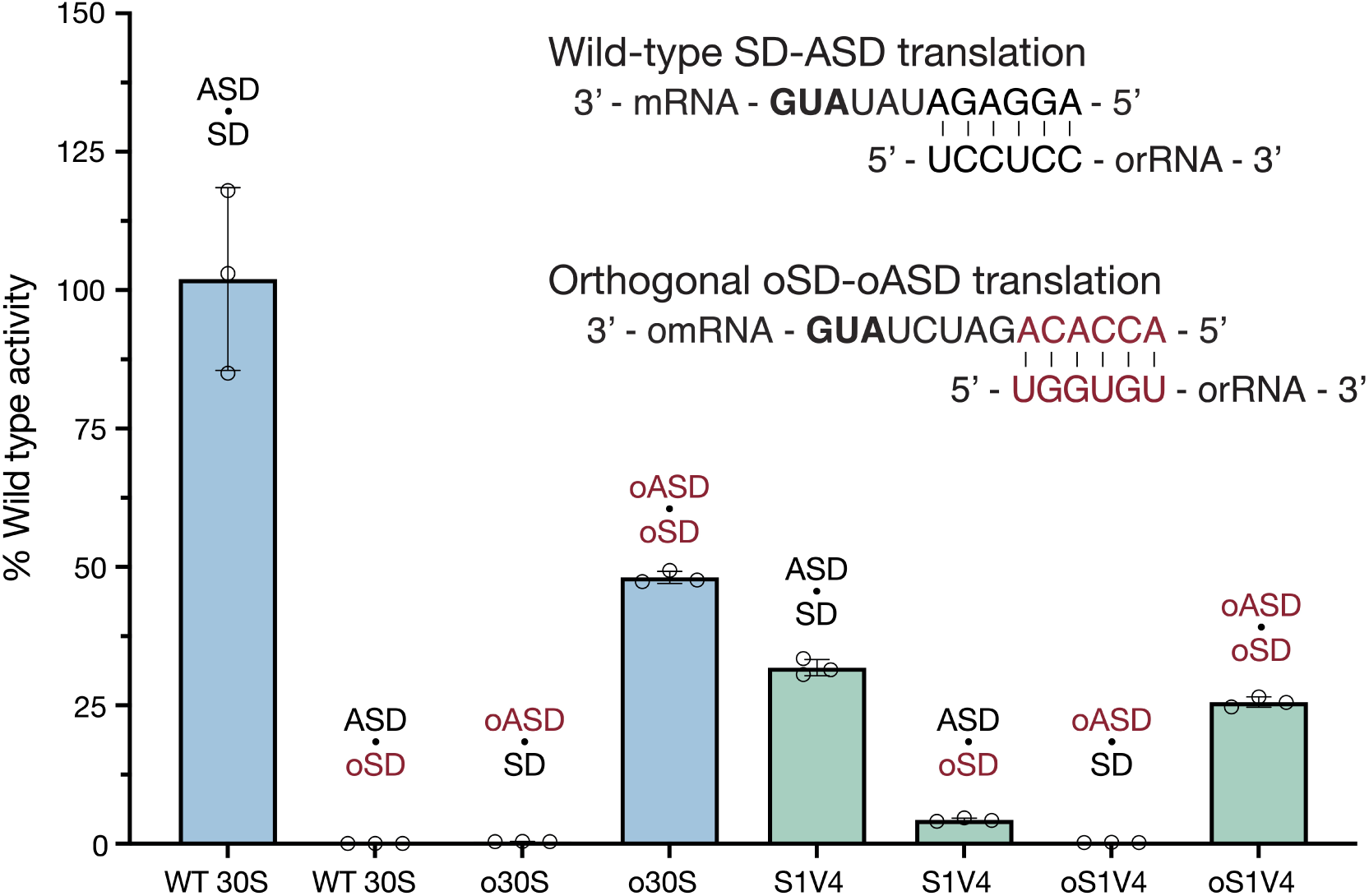
*In vitro* activity of 30S subunit S1V4. Activity of 30S ribosomal subunits with wild-type or orthogonal ribosome binding site elements in an *in vitro* nanoluciferase translation assay. Normalized luminescence of wild-type 30S subunits (blue) with a wild-type ASD sequence and oASD sequence with corresponding nanoluciferase reporters (black and red). The activity of 30S subunit S1V4 is shown in green with a wild-type ASD sequence and oASD sequence with corresponding nanoluciferase reporters (black and red). The activity is normalized to wild-type ribosomes. Error bars indicate the standard deviation for n=3 measurements.

Although preventing bS1 binding to the 30S subunit did not improve ribosome orthogonality *in vitro*, under these conditions there is no competition between mRNAs for ribosome binding as would occur in cells.^38^ We therefore sought to understand how preventing ribosomal protein bS1 binding to the 30S subunit would affect the orthogonality of ribosomes *in vivo*. To accomplish this, we developed a two-plasmid system based on previously designed reporters.^37^ In this system, GFP is encoded on a plasmid under the control of an arabinose inducible promoter, while the ribosomal RNA operon is encoded on a plasmid under control of an IPTG inducible promoter. In this system, wild-type GFP is translated from an mRNA harboring a wild-type SD, using ribosomes carrying the wild-type ASD (Figure 4A). In the orthogonal case, oGFP is translated from an mRNA harboring an orthogonal SD and should only be translated by ribosomes with an orthogonal ASD (Figure 4A). With the set of mRNA and ribosome plasmids harboring all combinations of SD, ASD, wild-type 16S, and S1V4 16S sequences, we co-transformed eight different variations of ribosome and GFP plasmids to test for ribosome orthogonality.

**Figure 4.**
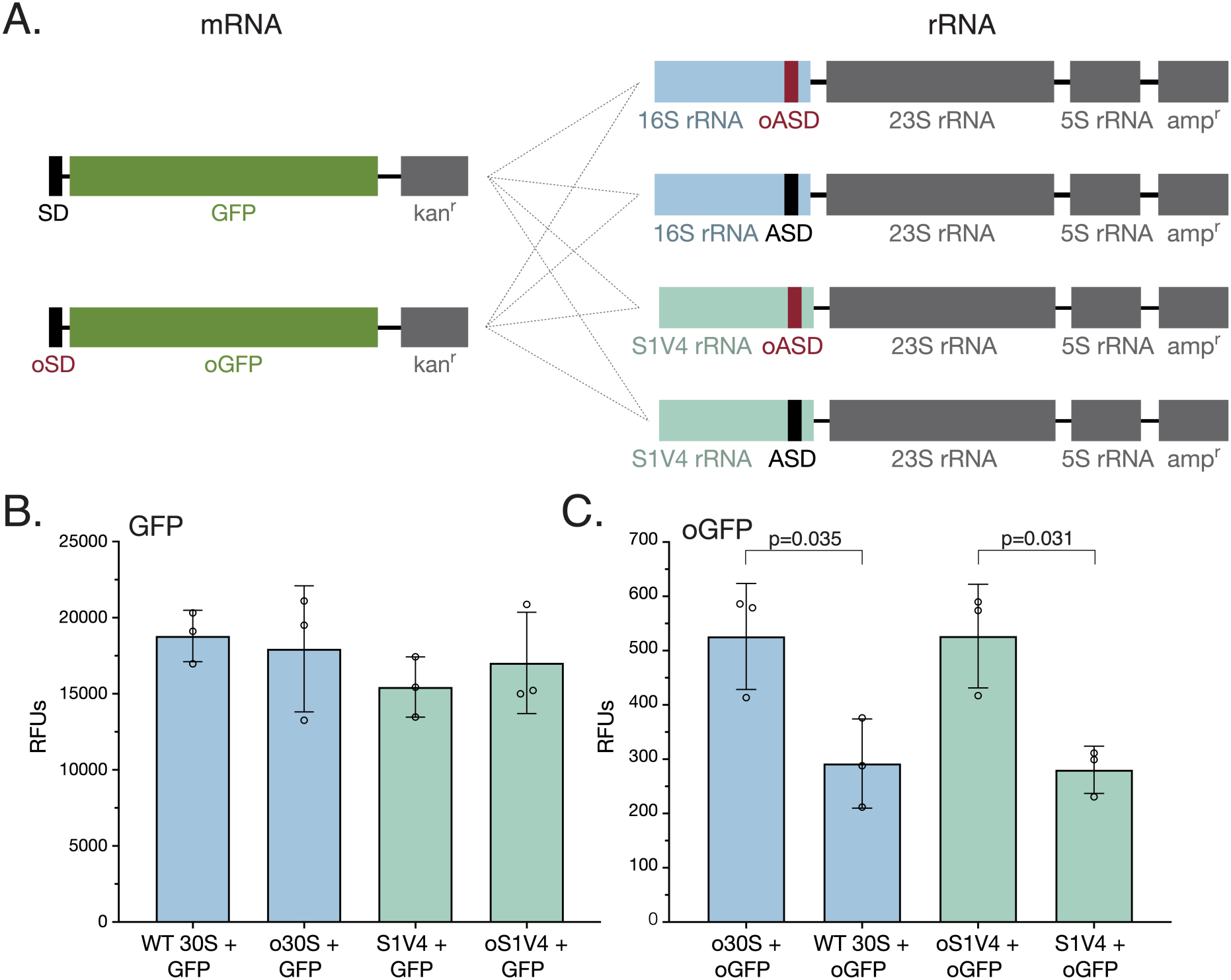
*In vivo* activity of S1V4 30S subunits. **A.** Two-plasmid reporter system with an arabinose inducible promoter upstream of GFP or oGFP, and an IPTG inducible promoter upstream of wild-type or S1V4 16S rRNA sequence, either with an ASD or oASD at the 3′ end. All 8 combinations used in reporter assays are indicated by connecting lines. **B.** *In vivo* translation of wild-type GFP by wild-type 30S (blue), o30S (blue), S1V4 (green), and oS1V4 (green). Error bars indicate standard deviation for n=3, p-values > 0.29. **C.** *In vivo* translation of orthogonal GFP by o30S (blue), wild-type 30S (blue), oS1V4 (green), and S1V4 (green). Error bars indicate standard deviation for n=3 measurements. Measurements reported in relative fluorescence units (RFUs).

We measured the fluorescence of GFP four hours after inducing GFP and ribosomal RNA expression with arabinose and IPTG, respectively. We also assessed the level of exogenous 16S rRNA expression using RT-PCR (Supplementary Figure 7). Cells from both wild-type 30S subunits and S1V4 30S subunits harboring the wild-type ASD and expressing the wild-type SD mRNA had comparable levels of fluorescence, which likely derives mostly from endogenous ribosomes given the level of exogenous 16S rRNA expression (Figure 4B, Supplementary Figure 7). We also observe high levels of fluorescence from cells expressing GFP from the mRNA with a wild-type SD (wild-type GFP) and o30S and oS1V4 30S, again likely due to the mRNA being translated by endogenous ribosomes. Interestingly, in the case of cells expressing oGFP mRNA, the amount of translation by ribosomes formed from oS1V4 30S subunits is comparable to that of cells expressing o30S subunits (Figure 4C), consistent with the *in vitro* translation results. The negligible amount of GFP cross-translation by wild-type ASD containing ribosomes of oSD mRNA is also comparable between wild-type and S1V4 30S subunits (Figure 4C). Taken together, these results indicate that the oSD and oASD sequences function similarly, independent of whether bS1 is able to bind to the 30S subunit or not.

## DISCUSSION

While orthogonal ribosomes play an essential role in efforts to expand the genetic code, they are limited by Shine-Dalgarno independent translation. This has been shown by single molecule tracking of orthogonal ribosomes in live cells, which revealed that 30S subunits, regardless of having an orthogonal ASD, still translate endogenous mRNAs.^13^ This is further supported by ribosome profiling experiments, using ribosomes with altered SD sequences, where it was shown that the SD sequence does not dictate the ability of the ribosome to translate endogenous mRNA.^14^ One possible reason for the cross-reactivity of endogenous mRNAs with orthogonal ribosomes is the role ribosomal protein bS1 plays in translation initiation. It has been hypothesized that bS1, which harbors nucleic acid binding domains, shuttles mRNAs to the ribosome by binding the 5′ region of the mRNA.^18,17^ This could also occur with orthogonal ribosomes, preventing full orthogonality of the translation systems by allowing orthogonal ribosomes to translate endogenous proteins.

Here, we tested the hypothesis that preventing bS1 from binding to the *E. coli* 30S ribosomal subunit would enable orthogonal ribosomes to be more dependent on the mRNA SD sequence. Since bS1 is essential in *E. coli*,^14^ we devised a strategy that would leave endogenous bS1 and endogenous ribosomes intact, yet allow us to test the role of bS1 binding to orthogonal ribosomes. In structures of the 30S ribosomal subunit, protein bS1 binds predominantly to ribosomal protein uS2, and extends in the direction of 16S rRNA helix h26. We took inspiration from RNA nanostructures and identified RNA k-turns as promising candidates to insert into h26 to sterically block bS1 from binding uS2.^35^ We engineered 16S rRNA using a prototypical k-turn in four different extensions of helix h26. Using a combination of biochemistry and cryo-EM, we were able to identify one variant (S1V4) that prevented bS1 binding to the 30S subunit while remaining translationally active, and without substantially perturbing ribosome structure, i.e. uS2 binding.

We assessed the orthogonality of ribosomes harboring S1V4 30S subunits using both *in vitro* and *in vivo* translation reactions^37^, and found that ribosome orthogonality is not improved by preventing bS1 binding to the 30S subunit. In fact, the *in vitro* reactions, in which no mRNA competition is involved, suggest preventing bS1 binding to the 30S subunit might actually decrease SD-dependent orthogonality. This leads us to think that other characteristics of mRNA may have an impact on SD independent translation, independent of the role of bS1. For example, recent studies have shown that adenosines at positions -3 and -6 relative to the start codon, increase translation efficiency.^39^ This sequence pattern is prevalent in *E. coli*, and could enable orthogonal ribosomes to translate these endogenous mRNAs, regardless of the absence or presence of bS1. Based on a large screen of over 200,000 sequences, unstructured UTRs also increase translation efficiency,^26^ a result confirmed by ribosome profiling experiments.^14^ Thus, mRNAs with unstructured UTRs could also enable binding to orthogonal ribosomes. Taken together, these features of mRNA outside of the SD-ASD interaction and independent of bS1 binding to the 30S subunit suggest that further improvements to ribosome orthogonality may require a deeper understanding of translation initiation mechanisms in *E. coli*.

## ACCESSION CODES

Atomic coordinates for the 30S models have been deposited with the Protein Data Bank under accession codes 9DUK and 9DUL for S1V3 and S1V4, respectively. Cryo-EM maps have been deposited with the Electron Microscopy Data Bank under the accession codes EMD 47168 and EMD 47169 for S1V3 and S1V4 (70S maps), respectively.

## ACKNOWLEDGMENTS

We would like to thank Paul Tobias, Dan Toso, and Ravi Thakkar for their assistance in the cryo-EM data acquisition. We would also like to thank Amos Nissley and Chandrima Majumdar for their help with the cryo-EM data processing and modeling. In addition, we would like to thank Amos Nissley, Chandrima Majumda, and Yekaterina Shulgina for helpful comments on the manuscript. This work was funded by the NSF Center for Genetically Encoded Materials (C-GEM), CHE 2002182.

## AUTHOR CONTRIBUTIONS

The project was conceptualized by K.V.B. and J.H.D.C. Experiments were performed by K.V.B., R.A.B., and M.K. The manuscript was prepared by K.V.B., R.A.B, and J.H.D.C.

## NOTES

The authors declare no competing financial interests.

