## Supplementary Information for "Role of ribosomal protein bS1 in orthogonal mRNA start codon selection"

| | S1V3 | S1V4 |
|---|---|---|
| Model resolution (Å) | 2.56 | 1.80 |
| FSC threshold | 0.5 | 0.5 |
| Total non-hydrogen atoms | 51540 | 51586 |
| R.m.s. deviations from ideal values | | |
| Bond (Å) | 0.005 | 0.009 |
| Angle (°) | 0.816 | 1.22 |
| Molprobity score | 2.43 | 2.42 |
| Clash score | 14.14 | 13.61 |
| Romater outliers (%) | 3.56 | 3.66 |
| Ramachandran plot | | |
| Favored (%) | 94.92 | 94.92 |
| Allowed (%) | 5.03 | 5.03 |
| Outliers (%) | 0.04 | 0.04 |
| RNA validation | | |
| Angles outliers (%) | 0.043 | 0.061 |
| Sugar pucker outliers (%) | 0.265 | 0.027 |
| Average suiteness | 0.541 | 0.536 |

**Supplementary Table 1. Cryo-EM statistics.** Overall map and model statistics for the S1V3 and S1V4 structures.

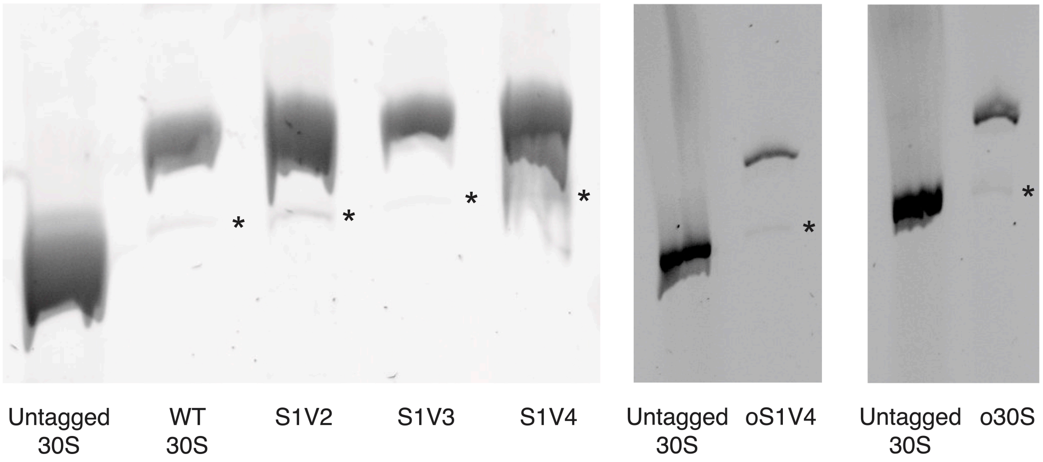


**Supplementary Figure 1. RT-PCR of ribosome mutants.** 10% TBE gel of cDNA from RT-PCR flanking the MS2 tag in the 16S rRNA. The lower band indicates wild-type 16S contamination, marked by asterisks. In all mutants, the tagged ribosomes have less than 3% wild-type contamination.

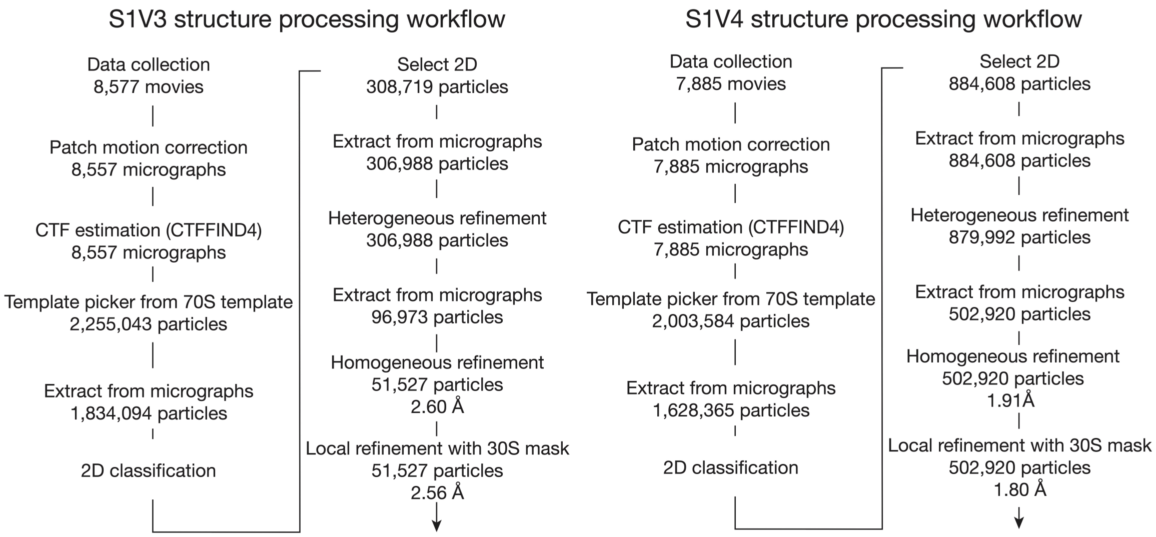


**Supplementary Figure 2. Cryo-EM processing workflow.** Cryo-EM data processing for both the S1V4 and S1V3 maps were done in CryoSPARC 4.[26] The number of movies, micrographs, or particles is listed at each step in the workflow. Resolutions are reported with an FSC threshold of 0.143.

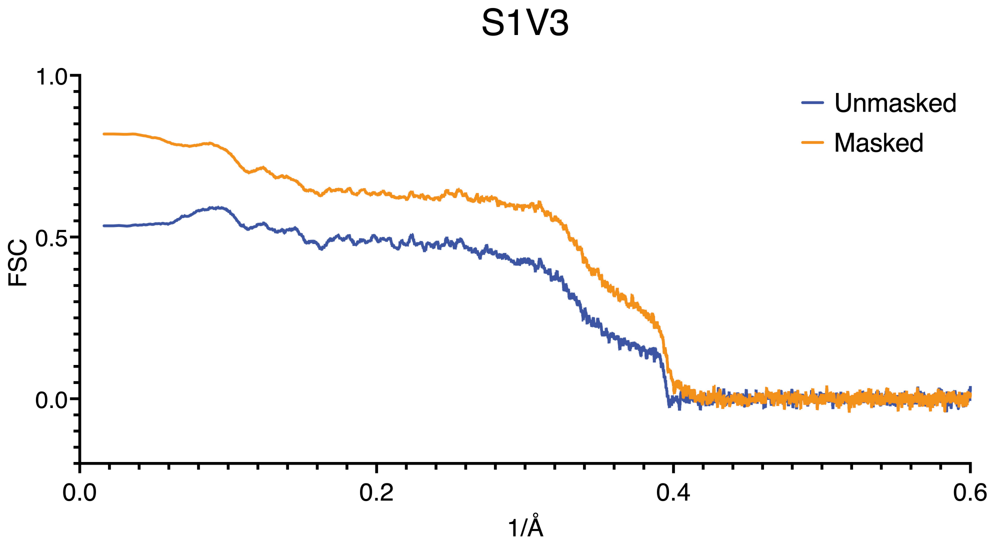


**Supplementary Figure 3. Fourier shell correlation (FSC) curves of the S1V3 map.** The FSC curve for the masked map (30S subunit only) is indicated in orange and the unmasked map (70S ribosome) is indicated in blue.

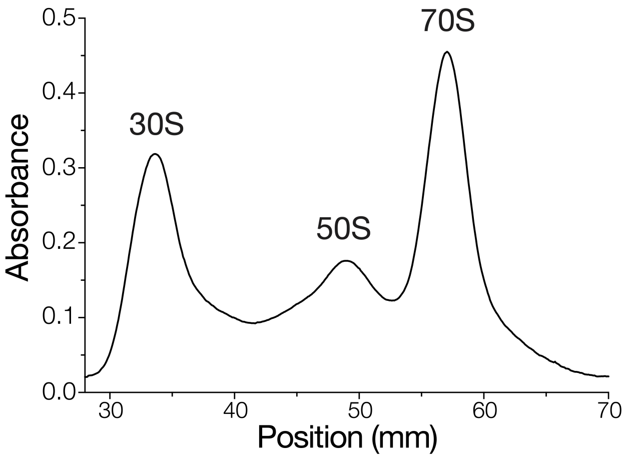


**Supplementary Figure 4. Reassociation of S1V4 30S subunits with wild-type 50S subunits.** 20-40% reassociation gradient of 1000 nM S1V4 30S subunits with 500 nM wild-type 50S subunits in 10 mM $MgCl_2$ and used for subsequent cryo-EM studies.

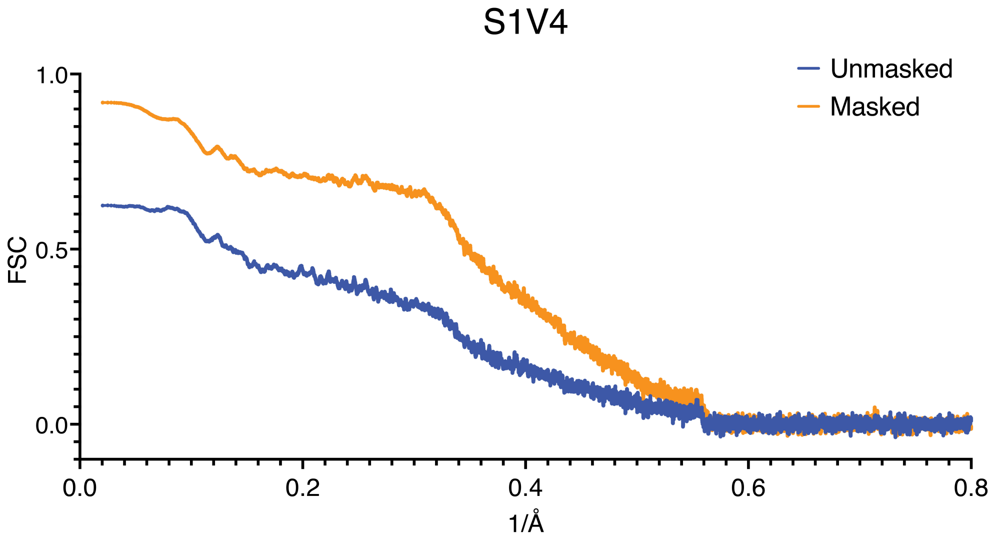


**Supplementary Figure 5. Fourier shell correlation (FSC) curves of the S1V4 map.** The FSC curves for the masked map (30S subunit only) is indicated in orange and the unmasked map (70S ribosome) is indicated in blue.

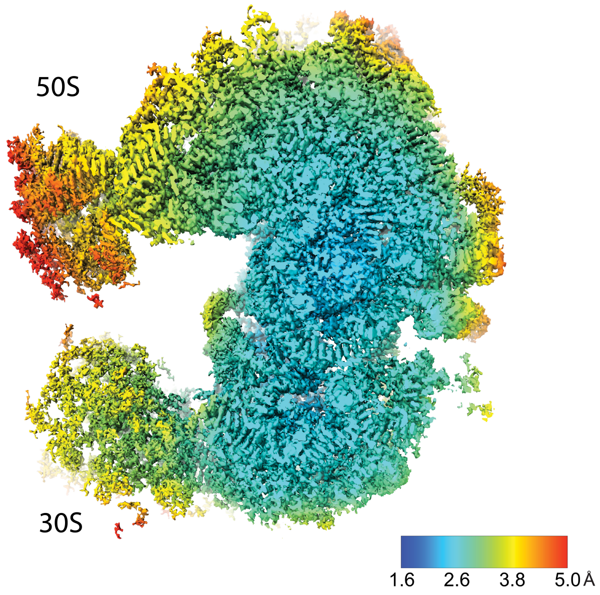


**Supplementary Figure 6. Local resolution of the S1V4 30S subunit cryo-EM map.** A cross-section of the local resolution was made in ChimeraX and the color key is reported in Ångstroms. Local resolution was measured using the Local Resolution estimation in Relion 4.0-beta-2.[29]

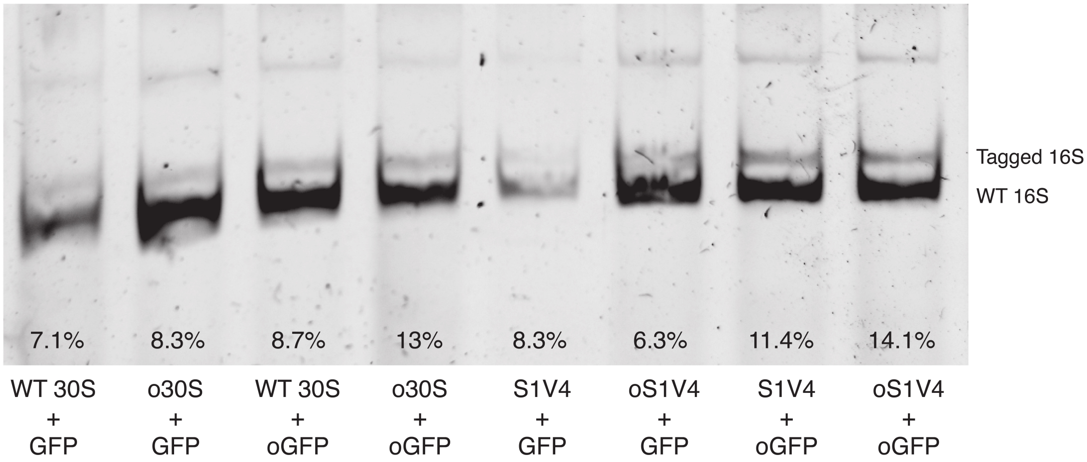


**Supplementary Figure 7. RT-PCR of mutant ribosomes**. 5% TBE gel of RT-PCR of mutant ribosomes used in *in vivo* experiments. RNA was extracted from cells four hours after induction with IPTG and arabinose. The lower band indicates wild-type 16S and the higher band is from the tagged 16S. The expected difference is 35 base pairs. Expression was quantified by integrating the gel bands, excluding the edges.
